# Lysosomal proteome and lipidome analyses of intestinal cells reveal the crucial role of bis(monoacylglycero)phosphate for autophagosome-lysosome fusion

**DOI:** 10.64898/2026.09.21.753291

**Authors:** Gianmarco Del Gallo, Zilei Chen, Anne Sanner, Robert Hardt, Fabrizio Merciai, Fabiola Salsano, Shroddha Bose, Michaela Schweizer, Diego L. Medina, Tobias Stauber, Jens Bosse, Eduardo M. Sommella, Dominic Winter, Sabrina Jabs, Thomas Braulke

**Author notes:** CONTACT: Thomas Braulke, mail; address: Department of Osteology & Biomechanics, Cell Biology of Rare Diseases, University Medical Center Hamburg-Eppendorf, Martinistr. 52, N27, 20246 Hamburg, Germany; Sabrina Jabs, mail; Institute of Clinical Molecular Biology, Christian-Albrechts-University Kiel & University Hospital Schleswig Holstein Campus Kiel, Rosalind-Franklin-Straße 12, 24105 Kiel, Germany.

## Abstract

The transport of about 70 enzymes to lysosomes depends on mannose 6-phosphate signals formed by GNPTAB. Editing of *Gnptab* in an intestinal mouse cell line revealed the loss of multiple lysosomal enzymes associated with the accumulation of sphingomyelins, ceramides, and cholesterol, and reduced levels of bis(monoacylglycero)phosphate (BMP) in lysosomal proteomes and lipidomes. By cross-correlation we identified two subsets of lysosomal lipid-modifying enzymes linked with mixed unsaturated or di-monosaturated BMP. Autophagy-related proteins functioning in early stages of autophagosome formation associated with neutral ceramides, whereas proteins involved in autophagosome-lysosome fusion correlated with negatively charged BMPs. Therefore, the high cholesterol and low BMP level of lysosomes might be causal for impaired autophagic flux in GNPTAB deficient cells. We propose a functional axis of three lysosomal proteins as potential target to improve the autophagic flux in cells with dysfunctional lysosomes and in a newly established intestinal organoid model suitable as novel experimental tool.

## Introduction

The *N*-acetylglucosamine-1-phosphotransferase (GNPTAB) plays a key role in the biogenesis of lysosomes. This Golgi-localized enzyme complex composed by each two α-, β- and γ-subunits, and its regulator protein LYSET, catalyzes the first step in the formation of mannose 6-phosphate (M6P) tags on high mannose type glycans of newly synthesized lysosomal enzymes. The M6P residues are recognized by M6P-specific receptors mediating the sorting of the enzymes in the *trans*-Golgi network and their delivery to lysosomes [1]. More than 70 luminal lysosomal enzymes and soluble accessory proteins are presently known to be involved in the low pH-dependent degradation of proteins, lipids, glycans, and nucleic acids reaching lysosomes via the biosynthetic, endocytic and autophagic pathways. The export of the degradation products to the cytosol and the regulation of lysosome ion homeostasis including lysosomal acidification require numerous transporters and channels localized in the lysosomal membrane [2]. In addition, lysosomes function in metabolic signaling, gene regulation, adaption to exogenous cues, communication and fusion with other organelles including autophagosomes via membrane contact sites and tethering factors and modify their intracellular positioning and mobility along microtubules [3–5].

Inherited biallelic mutations of the *GNPTAB* gene encoding the α- and β-subunits of GNPTAB associated with the complete loss of enzymatic activity cause a rare and fatal disease, mucolipidosis type II (MLII, also called I-cell disease; [6]). Clinically the patients suffer from progressive neurodegeneration, severe skeletal abnormalities (dysostosis multiplex), organomegaly and cardiorespiratory defects leading to death in their first decade of life [7]. Biochemically, the inability of mutant GNPTAB to tag soluble lysosomal proteins with M6P residues prevents their proper targeting to lysosomes and result in their hypersecretion into the extracellular space and bloodstream. However, analysis of cells and tissues of MLII patients or *Gnptab*-deficient mice has shown that small portions of distinct lysosomal enzymes can reach lysosomes via M6P-independent intracellular pathways or by secretion-recapture mechanisms [8–11]. These alternate transport routes are not sufficient to compensate for the loss of degradative capacity that leads to the accumulation of nondegraded macromolecules in lysosomes. Storage material in GNPTAB-deficient cells comprises all classes of biomolecules in a cell type-specific manner.

In the present study, we have edited the *Gnptab* gene in a non-tumorigenic, intestinal epithelial MODE-K cell line [12] and analyzed the proteome and lipidome of magnetite isolated lysosomal fractions. The reduction of 70% of luminal lysosomal enzymes in *Gnptab* KO cells was associated with a significant increase in 50 lipid species in particular sphingomyelin (SM) and ceramides, and reduced level of bis(monoacylglycero)phosphate (BMP) forms, in perinuclear lysosomes of enlarged size. Cross-correlation analysis between lysosomal enzymes and lipids suggested an interdependent acid sphingomyelinase/SMPD1, CLN5 and NPC2 network to regulate and sense three important classes of lipids, SMs, BMP and cholesterol. Using a similar approach, we correlated autophagy-related proteins with the lipidome of the purified lysosomes which identified two subgroups: The first subgroup comprised proteins involved in early stages of cargo binding to receptors, autophagosome biogenesis and selective autophagy that strongly correlated with ceramides and neutral glycosphingolipids, but not with BMP. The second subgroup comprised proteins functioning in later autophagy stages, such as autophagosome maturation and autophagosome-lysosome fusion which strongly correlated with BMP species. These data support our conclusion that cholesterol accumulation and reduced BMP levels are causal for the impaired autophagic flux in *Gnptab* KO cells. Finally, we have established 3D-intestinal organoid models from wildtype and *Gnptab* KI mice [13] and characterized them by proteomics, electron microscopy and lattice light sheet microscopy as a tool for future studies on multicellular systems with dysfunctional lysosomes and impaired autophagic flux.

## Results and Discussion

### Simultaneous loss of multiple lysosomal enzymes results in complex changes in the lysosomal lipidome

Recently Harper and colleagues [14] reported on the CRISPR-Cas9 targeting of more than 30 genes causing monogenic lysosomal storage diseases. Proteomic and lipidomic analysis of lysosomes purified from HeLa cell lines expressing endogenously HA-tagged TMEM192 for the immunoprecipitation revealed characteristic alterations in the lipid membrane composition correlating with the targeted lipid-metabolizing lysosomal proteins. In addition, these analyses provide novel insights into cellular pathways that depend on lysosomal homeostasis including autophagic and endocytic cargo delivery to lysosomes and alterations in the mitochondrial proteome. Here, we cross-correlated the lysosomal proteome and lipidome of the murine non-tumorigenic, intestinal epithelial MODE-K cell line deficient in GNPTAB which results in a complex dysfunction of lysosomes due to a simultaneous missorting of almost all luminal lysosomal proteins lacking M6P-tags [1]. We isolated magnetite-loaded lysosome fractions from parental wild type (WT) and *Gnptab* KO MODE-K cells followed by MS (Figure 1A; [10,15]). About 5200 proteins were identified. As expected, of the 60 detectable luminal lysosomal proteins, 42 were significantly reduced in their abundance in lysosomes of *Gnptab* KO cells including members of all classes of lysosomal hydrolases. In contrast, the majority of detectable lysosomal membrane proteins were not or weakly affected in mutant cells (Figure 1B; Table S1). These findings were validated for two lysosomal proteases, cathepsin D (CTSD) and Z (CTSZ) by western blotting, which showed reduced intracellular abundance, impaired proteolytic maturation and hypersecretion of inactive enzyme precursor forms into the medium (Figure 1C). Similar observations have been reported in various *GNPTAB* KO cell lines and patient fibroblasts [15–18].

**Figure 1.**
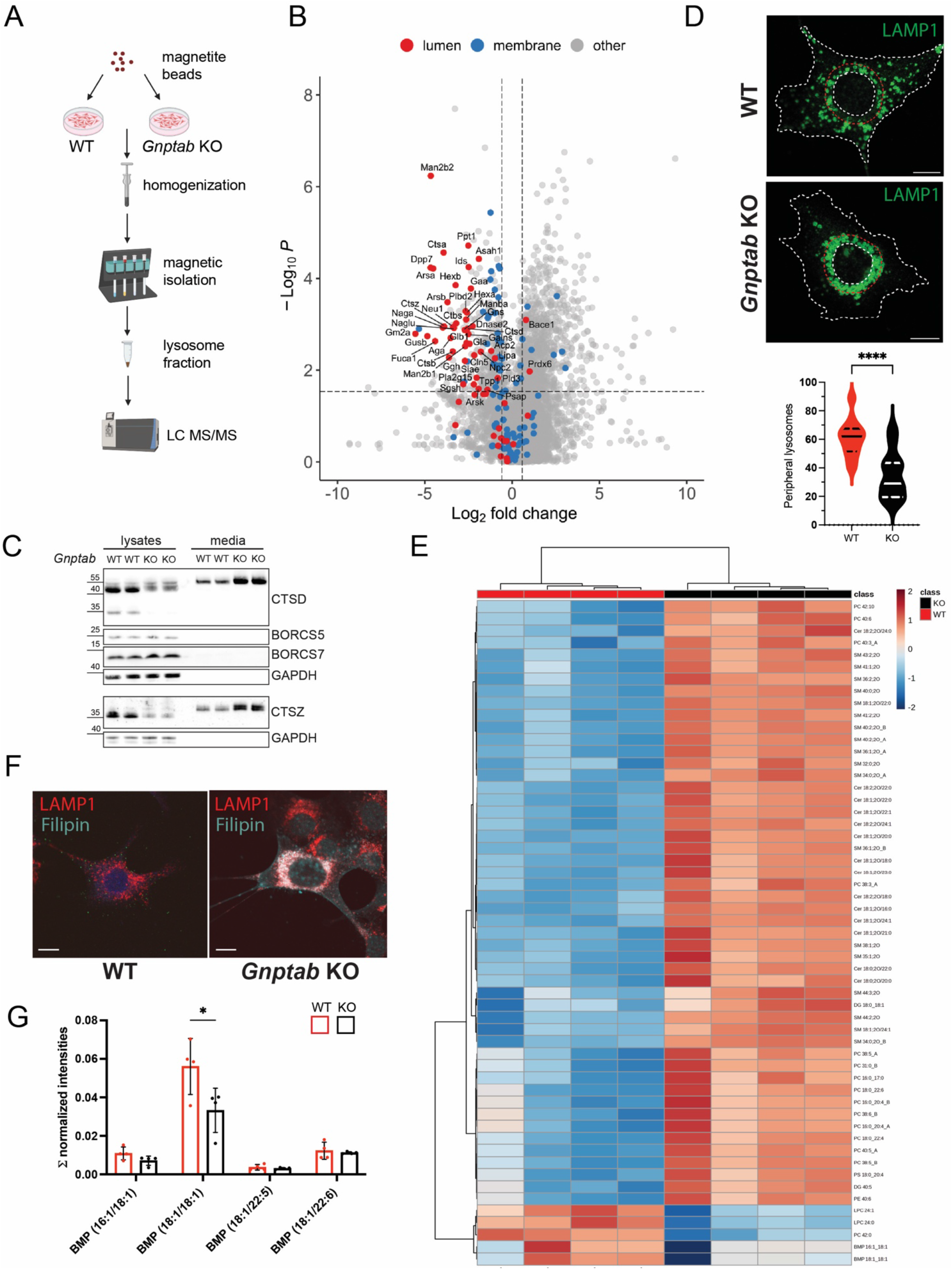
GNPTAB-deficiency leads to loss of lysosomal enzymes and changes in size and position of lysosomes in MODE-K cells. (**A**) Schematic workflow of magnetite-based isolation of lysosomes and LC-MS proteome analysis. Created in BioRender. Jabs, S. (2026) https://BioRender.com/fpbvo02 (**B**) Volcano plot of soluble luminal poteins (blue dots) and membrane proteins (blue dots) of isolated lysosomes from WT and *Gnpt*ab KO MODE-K cells (WT:*Gnpt*ab KO ratio) identified by LC-MS (mean SD, *n=*3) (**C**) LAMP1 (green) staining of WT and *Gnpt*ab KO cells and analyzed perinuclear areas are shown. Scale bar, 10 μm. The number of peripheral lysosomes in WT and *Gnpabt* KO MODE-K cells (n=25 cells per genotype) is shown. Unpaired t-test, **** *p*-value <0.0001. (**D**) Immunoblot analysis of indicated lysosomal proteins in lysates and conditioned media of WT and *Gnptab* KO MODE-K cells. GAPDH is used as loading control. (**E**) Heat map of differentially expressed lipids in isolated lysosome fractions of parental (WT) and GNPTAB*-*deficient MODE-K cells (normalized concentration; n=4). (**F**) Non-esterified cholesterol (visualized by Filipin; cyan) in lysosomes of WT and *Gnptab* KO cells co-stained with LAMP1 (red). Scale bars 10 μm. (**G**) BMP species in isolated lysosome fractions quantified as normalized concentration intensity (mean ± SD; unpaired student t-test; n=4 of each genotype; * *p*-value <0.05).

The missorting of newly synthesized lysosomal enzymes and their intracellular deficiency in *Gnptab* KO MODE-K cells resulted in an increased number of LAMP1-positive lysosomes enlarged in size and preferentially localized to the perinuclear region, whereas WT cells contained smaller lysosomes that were evenly distributed throughout the cytoplasm (Figure 1D). To examine whether the altered positioning of lysosomes was associated with changes in the abundance of proteins involved in lysosomal positioning and mobility, we analyzed subunits of BLOC-one-related complex, BORCS-5 and BORCS-7 [19] by western blotting. No significant changes in their abundance were observed consistent with the proteomic data which revealed that 22 out of 25 detected proteins involved in positioning and mobility including BORCS-5, -6 and -7 were not significantly altered (Table S1). These data might suggest that the molecular protein machinery regulating the lysosome trafficking are present but fail to function properly in *Gnptab* KO cells. Several studies report on alterations in positioning and size of lysosomes in cells defective in sphingolipid and cholesterol catabolism [20–23]. This prompted us to examine whether changes in lipid composition in the lysosomal membrane contribute to the altered distribution and size of lysosomes in *Gnptab* KO cells. We characterized the lipidomic landscape of magnetite-isolated lysosome fractions from WT and *Gnptab KO* MODE-K cells. Untargeted lipidomics profiling detected in total 339 lipid species covering 14 lipid classes (Table S2). Principal component analysis (PCA) displayed a genotype-dependent tight clustering of samples, indicating distinct changes in lipid composition of lysosomes between WT and *Gnptab* KO cells (Fig. S1A). Among the 55 significantly altered lipid species, 50 were increased and 5 reduced in *Gnptab* KO lysosomal fractions (Figure1E). The most significantly increased lipid classes were sphingomyelins (SMs) and ceramides (Cer; Fig. S1C) whereas lysophosphatidylcholines (LPC; Fig. S1C) and BMP species, in particular the most abundant BMP (18:1:18:1) form (Figure1G, Fig. S1B), were reduced in *Gnptab KO* lysosomes. BMP is synthesized in the acidic milieu by the lysosomal CLN5 protein [24]. In contrast to the almost complete loss of BMP in *CLN5*-KO HEK293T, *CLN5*-deficient iPSCs, and iNeurons, we have observed a partial reduction or no significant changes depending on the BMP species (Figure1G), suggesting that a fraction of newly synthesized CLN5 is transported to lysosomes and sufficient to support BMP synthesis in a cell type-specific and M6P-independent manner in MODE-K cells [9,10].

The observed massive accumulation of SMs is expected to be the result of reduced acid sphingomyelinase (ASM) activity, but the ASM protein (encoded by the *Smpd1* gene) was not significantly reduced in the lysosomal proteome of *Gnptab* KO MODE-K cells (Table S1). However, ASM/SMPD1 requires anionic lipids such as BMP for its recruitment to the intraluminal lysosomal membrane and for stimulation of enzymatic activity [25,26]. Therefore, the reduced BMP concentration in enlarged *Gnptab* KO lysosomes (Figure1E, G) due to the missorting of the lysosomal BMP synthase CLN5 may result in reduced lysosomal ASM/SMPD1 activity and thereby contribute to SM accumulation. Since BMP also stimulates the NPC2-mediated cholesterol egress from lysosomes [27] the loss of both NPC2 and CLN5 from *Gnptab* KO lysosomes as a consequence of defective M6P tagging (Figure1B) lead to low BMP level and secondarily cholesterol accumulation. This notion was corroborated by the strong filipin staining of non-esterified cholesterol in LAMP1-positive lysosomes of *Gnptab* KO MODE-K cells (Figure1F). In addition to CLN5, we observed a strong cross-correlation of several luminal lysosomal enzymes (including NAAA, PPT1, PPT2, HEXB, SMPD1, and GLA) and lysosomal membrane transporter (including SLC44A2, ABCD4, ABCA3, NPC1 and SCARB2/LIMP2) with BMP (18:1/22:5) and BMP (18:1/22:6) species (Fig.S2). Moreover, FUCA1, GLB1, NPC2, PLA2G15 and SPNS1 were strongly correlated with BMP (18:1/18:1) and BMP (16:1/18:1) (Fig.S2). The functional significance of these correlations remains to be investigated. Together, these findings suggest that ASM/SMPD1, CLN5 and NPC2 form an interdependent axis that regulates and senses three important classes of lipids, SMs, BMP and cholesterol, in lysosomal membranes, which is severely disturbed by defective M6P-tagging of luminal lysosomal proteins. In analogy, in WT cells the newly synthesized lysosomal enzymes and accessory luminal proteins carrying M6P tags need to be immediately dephosphorylated by two lysosomal phosphatases, ACP2 and ACP5, upon arrival in lysosomes. The removal of M6P has important functional consequences for several lysosomal enzymes. In particular, enzymes and accessory proteins involved in lipid metabolism including ASM/SMPD1, CLN5, NPC2, PPT1, GLB1, PLA2G15, have predicted isoelectric points between 6.0 and 8.7 disregarding signal peptides and posttranslational modifications such as glycosylation or phosphorylation. In cells lacking both ACP2 and ACP5 all other soluble lysosomal enzymes are properly targeted to lysosomes, but the M6P tags remain attached and prevent their efficient protonation at the luminal pH of approximately 4.5. We propose that the electrostatic interaction with intralysosomal membranes enriched with negatively charged lipids such as BMP, is impaired and subsequently the NPC2-mediated egress of cholesterol [28].

Whether the SMs accumulate to toxic concentrations resulting in lysosomal membrane permeabilization [29] remains unknown. The accumulation of Cer in *Gnptab* KO lysosomes most likely resulted from the deficiency of acid ceramidase (ASAH1) and the sphingolipid activator protein saposin D generated from prosaposin (PSAP) by CTSD and CTSB [30]. The abundance of ASAH1, PSAP, CTSD, and CTSB lacking the M6P targeting signals was significantly reduced in *Gnptab KO* lysosomes (Figure1B, D).

### Lysosomal lipid landscape in Gnptab KO cells impairs autophagy

Macroautophagy (hereafter referred to as autophagy) is a conserved pathway by which cellular material, including protein aggregates, damaged organelles or intracellular pathogens, is sequestered to autophagosomes that subsequently fuse with lysosomes for cargo degradation [5,31]. Several steps of the autophagic-lysosome pathway are mediated by an expanding molecular machinery of proteins. In the isolated lysosome fraction from *Gnptab* KO cells we found 16 autophagy proteins with increased abundance and 5 proteins with lower abundance compared with WT lysosomes (Figure2A; Table S1). At steady state, non-starved *Gnptab* KO MODE-K cells exhibited a marked increase in LC3-B immunoreactive bands by western blotting (Figure2B). Consistently, immunofluorescence microscopy showed weak and small cytoplasmic LC3-positive puncta in WT cells, whereas *Gnptab* KO MODE-K cells contained prominent LC3-stained autophagosomal structures (Figure2C). To assess the dynamics of autophagosome maturation and cargo degradation (autophagic flux), we expressed the tandem fluorescent reporter RFP-GFP-LC3 construct [32] in WT and *Gnptab* KO MODE-K cells and monitored the pH-dependent reduction in GFP fluorescence relation to the unchanged fluorescence of the acid-stable RFP fluorescence. The higher GFP:RFP ratio in *Gnptab* KO cells indicate a block in the autophagic flux (Figure2D, E). To exclude impaired lysosomal acidification of lysosomes in *Gnptab* KO cells as the cause for the increased GFP:RFP ratio in *Gnptab* KO cells, we performed ratiometric pH measurements using dextran-coupled pH-sensitive Oregon Green® 488. Lysosomal pH was unchanged between the two genotypes, measuring 4.4 ± 0.1 in both cell lines (Figure2F). Together, these data demonstrate that the autophagic flux in *Gnptab* KO MODE-K cells is strongly impaired.

**Figure 2.**
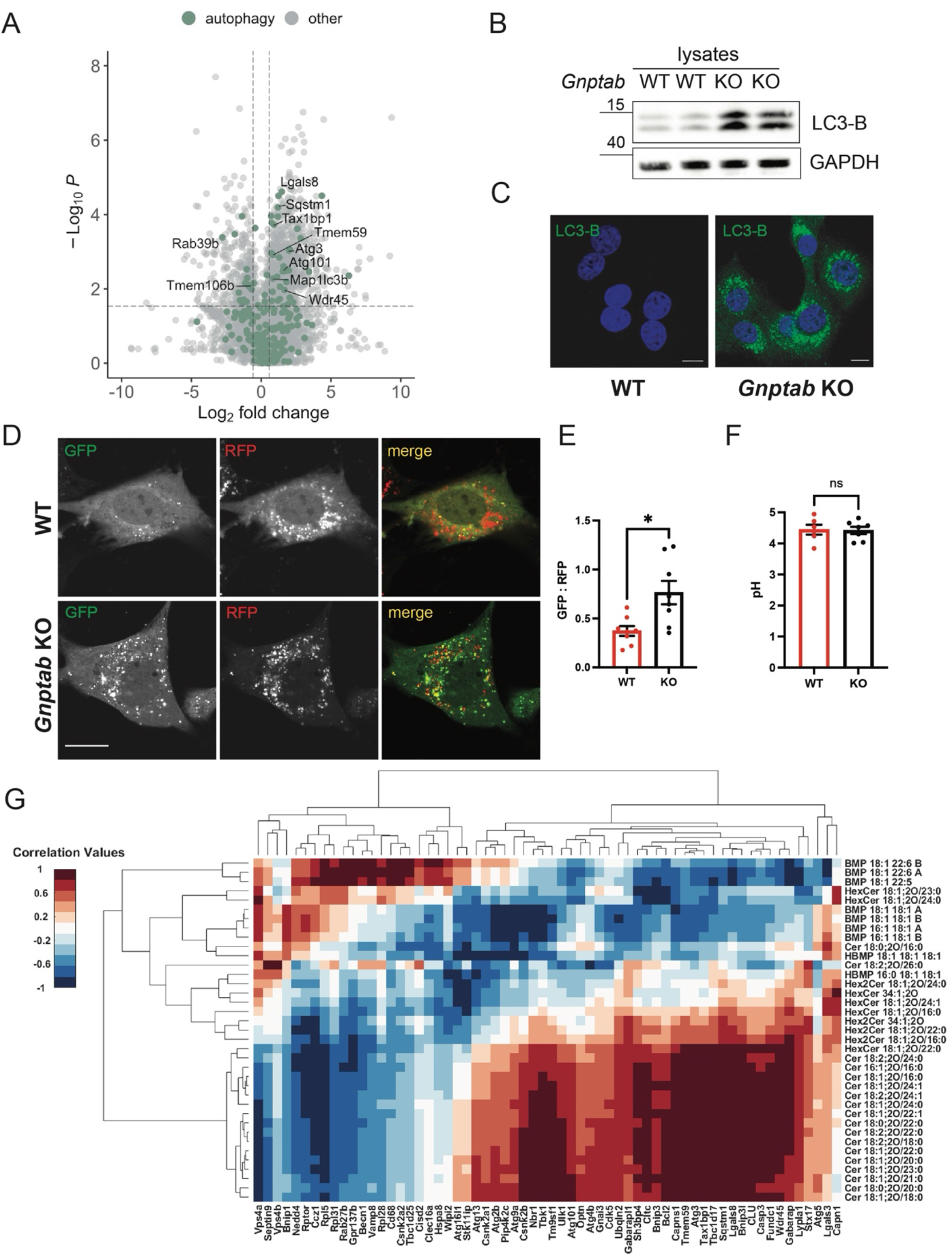
Autophagy-related proteins, lysosomal lipid correlation, and block of autophagic flux in Gnptab-KO MODE-K cells. **(A)** Volcano plot of autophagy-related proteins in isolated magnetite-lysosomal fractions (wt: Gnptab-KO ratio). Log2-fold cutoff, 0.58; p-value cutoff q< 0.05. **(B, C)** Strong accumulation of endogenous LC3-B by immunoblotting, and immunofluorescence microscopy (green) and DAPI (blue); scale bar 10 µm. **(D)** Monitoring autophagic flux in WT and *Gnptab* KO MODE-K cells transiently expressing RFP-GFP-LC3 without treatment. Scale bar 10 µm. **(E)** The GFP:RFP fluorescence ratio was determined in n=8 cells of each genotype of two independent experiments. **(F)** Ratiometric imaging of lysosomal pH measurements in WT and Gnptab-KO cells using dextran-coupled pH-sensitive Oregon Green® 488. Mean ± S.E. measuring the lysosomes of at least 10 different cells per genotype; n=6. **(G)** Autophagy-related protein-lipid correlation (Letters “A” and “B” indicate possible isomers).

When we correlated autophagy-related proteins with the lipidome of the lysosomal fractions, we identified two protein subgroups (Figure2G). Several proteins of subgroup 1 (including VPS4A, VPS4B, SEPTIN9, BNIP1, NEDD4, RPTOR, CCZ1, RPL5, RPL31, RPL28, GPR137B, BECN1, VAMP8, CD68, CSNK2A2, TBC1D25, CISD2, CLEC16A, WIPI2, ATG16L1, STK11Ip) function at the interface between core initiation, autophagosome maturation, late-stage autophagosome-lysosome fusion, and ribophagy [33,34], This subgroup of autophagy proteins showed a strong positive correlation with BMP (18:1/22:6) and BMP (18:1/22:5) species and a weaker correlation with BMP (18:1/18:1), and BMP (16:1/18:1), but exhibited a negative correlation with many ceramide species. In contrast, subgroup 2 comprising about 40 autophagy-related proteins including several receptors OPTN, TAX1BP1, NBR1, BENIP3L, SQSTM1, and LGALS8, and correlated predominantly with ceramides, partially with glycosphingolipids, but not with BMP species (Figure2G). These proteins act in the early steps of core initiation, elongation, lipid conjugation, ubiquitin-based mitophagy, xenophagy and aggrephagy, sensing of organelle damage, and the crosstalk between autophagy and apoptosis [35–38]. Together, these protein-lipid correlations revealed distinct lipid signatures associated with different stages of the autophagy pathway. Neutral lipid (sphingolipid, and glycosphingolipid) fingerprints which were primarily associated with proteins involved in early steps of cargo binding to receptors and the biogenesis of autophagosomes (subgroup2), whereas negatively charged BMP species appeared to be essential for the late-stage autophagosome maturation and autophagosome-lysosome fusion machinery (subgroup 1). These findings suggest that both the cholesterol accumulation and the reduced BMP abundance in endo/lysosomal membranes of *Gnptab* KO MODE-K cells may be causal for the marked impairment of autophagic flux. The molecular mechanisms by which these lipids regulate autophagosome-lysosome fusion remain to be elucidated.

### Establishment of a 3D-multicellular Gnptab KI intestinal organoid model

In addition to the murine intestinal *Gnptab* KO MODE-K cells, we used our *Gnptab^c.3082insC^* knock-in (*Gnptab* KI) mice, the closest disease model with a common mutation of MLII patients [39–41] to establish a more physiologically relevant multicellular 3D-intestinal organoid model. Live-cell confocal imaging was performed with WT and *Gnptab* KI organoids using LysoTracker™ Deep Red to label acidic organelles and the green fluorescent Bodipy-labeled lactosylceramide (Bodipy-LacCer) to visualize membrane domains (Figure3A, B). In WT organoids, lysosomes appeared as small puncta distributed throughout the cytoplasm. In contrast, in *Gnptab* KI organoids lysosomes were enlarged in size and formed clusters which preferentially localized beneath the apical surface facing the organoid lumen. To achieve higher spatial resolution and enhance image quality we performed live-organoid imaging by lattice light sheet (LLS) microscopy. (Figure3C, D). The differences in size and distribution of lysosomes in WT and *Gnptab* KI organoids were clearly confirmed by the reconstruction of the lysosome network (Figure3C and D). Most notably, small sized scattered lysosomes could be observed throughout the cells of WT organoids which showed motility in all directions, whereas the loss of GNPTAB resulted in the localization of enlarged and immobile lysosomes towards the apical cell periphery (Figure. 3E). Ultrastructural analysis by electron microscopy (EM) was performed on 60 nm thin sections of WT and *Gnptab* KI organoids. Due to the thin section thickness the small lysosomes in WT enterocytes, which constitute the majority of intestinal cells, were rarely detected (Fig. S3). In contrast, enlarged lysosomes filled with membranous storage material could be observed in *Gnptab* KI enterocytes.

**Figure 3.**
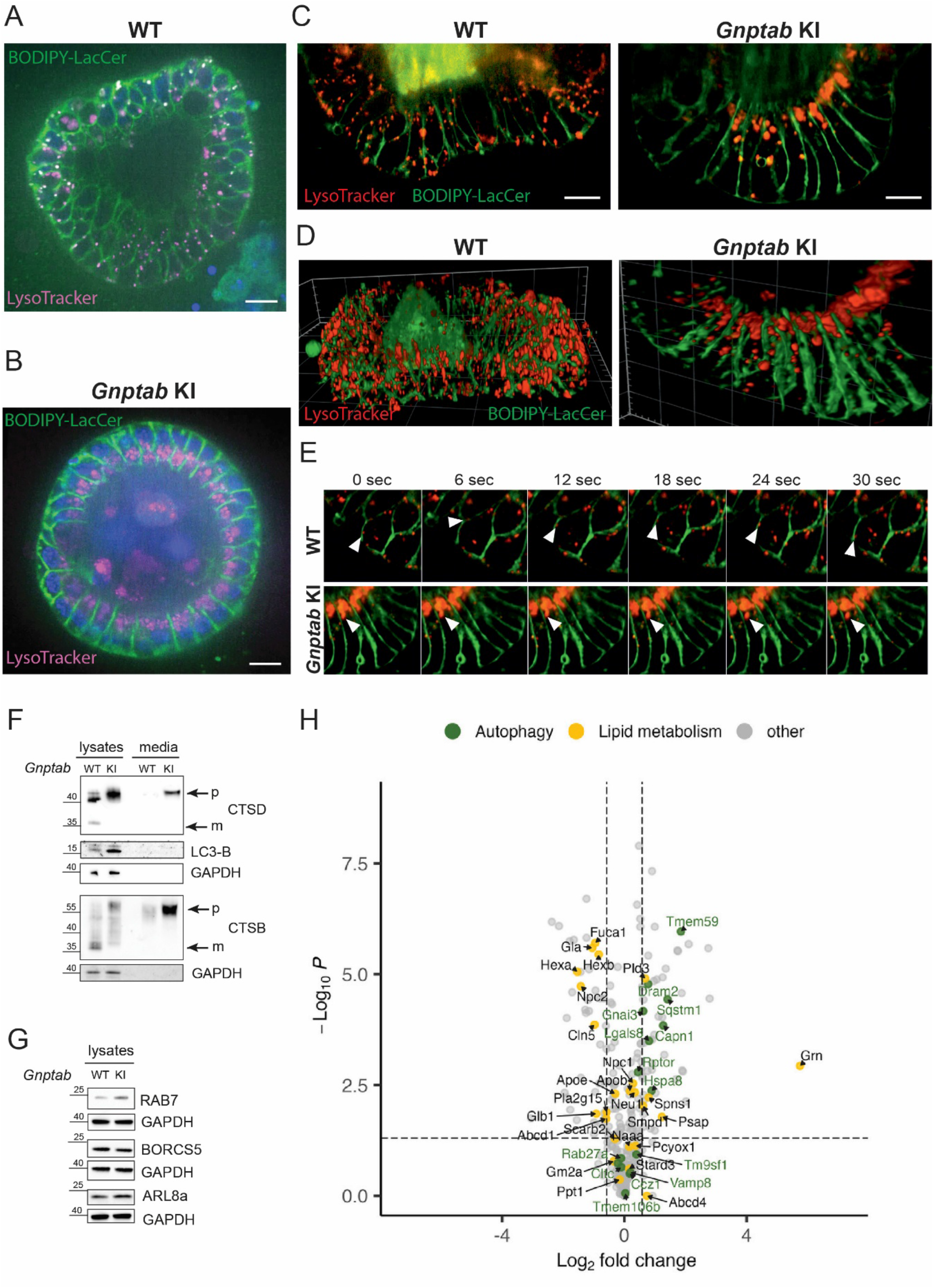
Positioning and mobility of lysosomes in intestinal organoids from WT and *Gnptab* KI mice. Lysosomes in intestinal organoids of WT (**A**) and *Gnptab* KI mice **(B)** were stained with LysoTracker Deep Red (purple), cell membranes with Bodipy-LacCer (green), and nuclei with Hoechst 33342 (blue). Representative snapshots from confocal live-imaging movies are shown. Scale bar 15 μm. (**C**) Lattice Light Sheet (LLS) live-cell imaging analysis of WT *Gnptab* KI organoids. Scale bar 10μm. (**D**) Lysosomal distribution reconstructions from LLS movies supported the different localization of lysosomes in WT and *Gnptab* KI organoids. (**E**) Representative snapshots from LLS movies showed the mobility of small lysosomes (white arrowheads) in cells of WT organoids tracked over 30 sec and the enlarged lysosomes of *Gnptab* KI organoids unable to move. (**F**) Volcano plot of autophagy-related proteins (green) and proteins involved in lipid degradation in lysosomes and egress (ocher) identified by targeted proteomics (WT: *Gnptab* KI ratio). Log2-fold cutoff, 0.58; p-value cutoff q< 0.05. (**G**) Western blot analysis of cathepsins (CTSD and CTSB) and LC3-B in lysates and conditioned media of WT and KI organoids. GAPDH served as loading control. (**H**) Western blotting of proteins involved in positioning of lysosomes from lysates of WT and KI organoids.

To characterize the intestinal *Gnptab* KI organoids biochemically, the lysosomal proteome was analysed by a targeted Mass Spectrometry approach. We applied a refined version of a previously established parallel reaction monitoring (PRM) assay for mouse lysosomal proteins [42], reliably detecting 321 proteins. Of those, 304 proteins were quantified with at least two values in one condition (WT or KO), including 28 soluble lysosomal enzymes and accessory proteins that were absent or significantly reduced in *Gnptab* KI organoids, most likely caused by the hypersecretion of enzymes lacking M6P tags. Only 5 lysosomal enzymes and co-factors were increased in their abundance in comparison with WT organoids: thiol protease cathepsin B (CTSB), the regulator of lysosomal proteolysis progranulin (GRN), the co-factor for sphingolipid degradation prosaposin (PSAP), the exonuclease digesting single-stranded DNA phospholipase D3 (PLD3), and ASM/ SMPD1 which converts sphingomyelin to ceramide (Table S3). Western blot analyses revealed that lysates of *Gnptab* KI organoids contained the inactive precursors but not the mature active forms of CTSD and CTSB (Figure3F). Increased amounts of the CTSD and CTSB precursors were secreted into the culture medium (Figure3F). Similar processing and localization defects of CTSD are detected in embryonic fibroblasts (MEFs) and isolated hepatocytes from *Gnptab* KI mice. In MEFs, the LDL receptor/LDL receptor-related protein 1 (LDLR/LRP1) mediate the secretion-recapture targeting route of CTSD and CTSB [10]. In the liver and in hepatocytes from *Gnptab* KI mice also mannose receptors which function as alternative M6P-independent receptors for the transport of non-phosphorylated lysosomal proteins are present in addition to the LDLR family [9,43]. The targeted proteomic dataset contained only a limited number of proteins involved in positioning and motility of lysosomes some of them were not detected in one replicate or entirely absent in one of the genotypes. Among the eight detected positioning proteins two, BORCS7 and ARL8a, were significantly reduced in *Gnptab* KI organoids, whereas the abundance of the others was not changed. Neither the lysosomal proteome of WT and mutant MODE-K cells revealed significant changes in the abundance of BORC7 and ARL8A (Table S1) nor western blotting of RAB7, BORCS5 and ARL8A (Figure. 3G). Together, these findings suggest that altered lysosomal positioning and reduced lysosomal motility in *Gnptab* KI organoids cannot be explained by major changes in the abundance of components of the known lysosomal trafficking machinery.

Finally, the abundance of proteins involved in lipid degradation and egress as well as in autophagy is summarized in Figure. 3H. Eight proteins, mostly involved in lipid degradation, were consistently reduced in *Gnptab* KI organoids and *Gnptab* KO MODE-K cells (PLA2G15, HEXA, HEXB, CLN5, NPC2, FUCA1, GLA, GLB1), and eight proteins involved in autophagy pathways were altered in both models (NCOA4, TMEM59, TMEM165, LDLR, ECE1, ATP11B, SQSTM1 and LGALS8).

In conclusion, these data demonstrate that our intestinal *Gnptab* KI organoid model recapitulates the biochemical and cell biological properties of the intestinal MODE-K cell line and provides an expanded, more physiological experimental system for investigating lysosomal homeostasis and related autophagy processes in a multicellular context. Integrating the lysosomal proteome and lipidome identified an ASM-CLN5-NPC2 axis that appears to sense and regulate the lysosomal membrane lipid composition, highlighting this pathway as a potential target for restoring autophagic flux in mucolipidosis II and other lysosomal disorders. Future studies using this organoid model will help to elucidate how alterations in lysosomal lipid composition affect lysosomal motility and how these changes subsequently impinge on autophagy.

## Materials and Methods

### Materials

Lipofectamine2000, Zeocin (Invitrogen); 4-nitrophenyl-N-acetyl-β-D-glucosaminide, human recombinant epidermal growth factor (EGF), N-acetyl-cysteine, Filipin complex , Protease Inhibitor Cocktail (PIC) powder, 2-(4-amidinophenyl)-1H-indole-6-carboxamidine (DAPI), poly-L-lysine, puromycin dihydrochloride (Sigma); dextran-coated magnetite DexoMAG 40 (Liquid Res. Ltd); LS magnetic columns (Miltenyi); Matrigel® Matrix Basement Membrane (phenol-red free), Corning® Cell Strainer 70µm (Sigma); Nitrocellulose membranes 0.45µm (Amersham Protran); Omnifix-F Solo 1ml (B. Braun); saponin (Fluka); µ-slide 8 Well Glass Bottom (Ibidi); JetPEI DNA Transfection Reagent (Polyplus); Aqua-Poly/Mount (Polysciences Inc); pGEM®-T Easy Vector (Promega); Triton X-100 (Roth);

Dulbecco’s Modified Eagle’s Medium (DMEM), Advanced DMEM/F12, fetal bovine serum (FBS), GlutaMAX, penicillin/streptomycin, Iscovés Modified Dulbeccós Medium (IMDM), Opti-MEM I were from Gibco.

#### Fluorophore probes

Bodipy Lactosylceramide complexed to BSA, LysoTracker Deep Red, Wheat Germ Agglutinin (WGA) Alexa Fluor555 conjugate (Invitrogen); WGA CF488A conjugate (Biotium); Filipin complex ready-made solution 5 mg/mL, 2-(4-amidinophenyl)-1H-indole-6-carboxamidine (DAPI) (both Sigma-Aldrich).

LC–MS grade water, acetonitrile (ACN), methanol (CH_3_OH), isopropanol (IPA), chloroform (CHCl_3_), methyl tert-butyl ether (MTBE), LC-MS grade additives formic acid (HCOOH) and ammonium formate (HCOONH_4_) were purchased from VWR (Milan, Italy). Deuterium labeled standards (15:0-18:1-d7-PC, 18:1-d7 Lyso PC, 15:0-18:1(d7) PE, 18:1(d7) Lyso PE, 15:0-18:1(d7) PG (Na Salt), 15:0-18:1(d7) PI (NH_4_ Salt), 5:0-18:1(d7) PS (Na Salt), 15:0-18:1(d7)-15:0 TAG, 15:0-18:1(d7) DAG, 18:1(d7) Chol Ester, d18:1-18:1(d9) SM, C15 Ceramide-d7) and authentic lipid standard mixtures (LightSPLASH®) were purchased by Avanti Polar Lipids (Alabaster, AL, U.S.A). Unless stated otherwise other reagents were all purchased by Merck.

### Cell culture and transfection

Murine duodenal epithelial cell clone K (MODE-K) cells [12] were maintained in DMEM supplemented with 10% heat-inactivated FBS, 25 mM HEPES pH 7.2, GlutaMAX™- and 1% penicillin/streptomycin. For transient transfection cells were seeded on poly-L-Lysine pre-coated coverslips or MatTek glass bottom dishes, and transiently transfected next day with cDNA constructs (0.5 µg) for 24 h using Lipofectamine 2000 according to the manufacturerś instructions. R-Spondin 1 and Noggin overexpressing HEK293 cells (kindly provided by Dr. Calvin Kuo (Stanford University) were cultured in DMEM containing 10% FBS and either 10 µg/mL puromycin (Noggin) or 0.3 mg/ml zeocin (R-Spondin 1). Conditioned media from R-Spondin 1 and Noggin-expressing cells were prepared in a large scale by mixing 4 batches of each ∼ 125 mL as described [44].

### Generation of Gnptab knockout MODE-K cells using CRISPR/Cas9

sgRNAs were designed using the SYNTHEGO sgRNA design tool (https://design.synthego.com/<u>#/</u>). DNA oligos containing sgRNA sequences for exon 3 (for CACCGCATGGGCAGACAGAGCCTA; rev AAACTAGGCTCTGTCTGCCCATGC) were annealed and ligated into BbsI-digested pX459 vector. MODE-K cells were transfected with pX459 constructs using JetPEI transfection reagent and selected with puromycin. Four days later, single clones were seeded into a 96-well plates, followed by expansion of 40 clones which were tested for missorted β-hexosaminidase by activity measurement [15] in 24-hour conditioned media, and western blotting for the loss of intracellular cathepsin B. Two potential *Gnptab KO* clones were genotyped by isolating genomic DNA followed by amplification (primers: for ACTCTATAGACAAGGCTGTCCTC; rev GTGCATCAGTTGTGGGTTACT), subcloning into pGEM T Easy Vector (Promega), and sequencing. Amplicon sequences were aligned to the reference sequence (GRCm39 assembly) using the BLAST analysis tool demonstrating deletions resulting in the premature termination in exon 3.

### Immunofluorescence microscopy

MODE-K cells seeded on 12 mm coverslips pre-coated with 0.1 mg/mL Poly-L-Lysine and cultivated for 48 h at 37 °C, 5% CO2. Cells were fixed in 4% PFA for 20 min at room temperature and incubated with 50 mM NH4Cl for 10 min at room temperature. Cells were permeabilized in permeabilization buffer (PBS containing 0.2% saponin and 3% BSA) for 10 min at room temperature, incubated with the primary antibody in permeabilization buffer at the indicated dilution overnight at 4 °C and with the appropriate Alexa fluorophore (coupled secondary antibody (1:1,000) for 2 h at room temperature. DAPI (1:2000) in permeabilization buffer was applied for 10 min at room temperature. Antibodies were: rabbit anti LC3B (1:100;Cell SignalingTechnologies); rat anti LAMP1 1DB4 (1:1000, DHSB). Coverslips were mounted on slides using Aqua-Poly/Mount and imaged using a using a confocal Leica TCS SP8 X microscope supported by a Leica LAS X software and analyzed with ImageJ.

For Filipin staining, cells were cultivated in complete DMEM for 24 h at 37 °C and 5% CO2. Cells were fixed and quenched as above and permeabilized with PBS containing 0.05% Triton X-100 for 10 min at room temperature. The staining was performed using 25 μg/ml of neutral polyene Filipin isolated from Streptomyces filipinensis for 45 min at room temperature in the dark, followed by dilutions of primary (overnight 4 °C) and secondary (2 h at room temperature) antibodies incubations in PBS (Kwiatkowska et al., 2014). After three washes in PBS, coverslips were mounted on glass slides with Aqua-Poly/Mount. Filipin was imaged with the 405 nm laser of a confocal Leica TCS SP8 X equipped with Leica LAS X software and analyzed with ImageJ.

### Mice

Heterozygous *Gnptab^c.3082insC^*(*Gnptab* KI) mice in C57Bl/6 or 129/SvJ genetic background were inbred to yield wild-type (WT) and homozygous *Gnptab* KI in a mixed background and genotyped for the WT and mutant alleles by PCR [13].The mice were housed in a pathogen-free animal facility at the University Medical Center Hamburg-Eppendorf, and experimental procedures were performed according to the institutional and ethical guidelines.

### Murine intestinal organoid culture

Intestinal crypts were isolated from mouse small intestines and cultivated according to the basic protocol by Mahe et al. [45] with slight modifications: The sedimented crypts (300 – 400) were resuspended in 50 µL Matrigel mixed with organoid growth medium (Basal Gut Medium (Advanced DMEM/F12 supplemented with HEPES, GlutaMAX, Penicillin/Streptomycin and 2 mM N-acetyl-cysteine) with 50 ng/mL EGF and conditioned media of R-Spondin 1 and Noggin in a ratio of 7:2:1), and placed in a 24-well plate. After incubation of the plates for 20 minutes at 37 °C to allow polymerization of Matrigel, 500 µL/well of organoid growth medium were added. Every 3 days the medium was replaced, and the organoids were passaged every 7 days and split in a 1:3 ratio.

### Organoid preparation for proteomic analysis

After 7 days in culture WT and *Gnptab* KI organoids were collected in ice-cold DPBS as described above, followed by centrifugation at 400 xg for 10 min at 4 °C [45]. The supernatant was removed and the organoid pellets were frozen. In total three replicates of both WT and *Gnptab* KI organoids were prepared. Each replicate contained organoids pooled from 7 wells, with an average of 15 organoids per well.

### Organoids live-cell confocal imaging

Organoids were passaged and seeded in an Ibidi µ-slide 8 Well during the weekly passaging. After 48h in culture, after 48 h in culture the organoids were incubated with 5 μM Bodipy Lactosylceramide complexed to BSA (1:100) in HBSS for 30 min at 4 °C, followed by three washes with ice-cold Basal Gut medium, and 50 nM LysoTracker Deep Red in pre-warmed Basal Gut medium for 30 min at 37 °C. After a quick wash in the same medium, organoids were imaged using a confocal Leica TCS SP8 X microscope equipped with Leica LAS X software and images analyzed with ImageJ.

### Lattice Light-Sheet Microscopy of organoids

Organoids were prepared and seeded in an Ibidi μ-slide 8 Well during the weekly passaging. After 48 h in culture the organoids were incubated with 5 μM Bodipy Lactosylceramide complexed to BSA (1:100) in HBSS for 30 min at 4 °C, followed by three washes with ice-cold Basal Gut medium, and 50 nM LysoTracker Deep Red in pre-warmed Basal Gut medium for 30 min at 37 °C. After a quick wash in the same medium, organoids were imaged using ZEISS Lattice Lightsheet 7 microscope equipped with Zen 3.2 software.

### Ultrastructural analysis of murine intestinal organoids

After 7 days in culture matrigel containing WT and *Gnptab* KI intestinal organoids were resuspended several times by pipetting in ice-cold PBS to break the matrigel droplets followed by centrifugation for 10 min at 400 xg [45]. The intestinal organoid pellets were fixed in a mixture of 4% paraformaldehyde and 1% glutaraldehyde (Science Services, Germany) in 0.1 M phosphate buffer at 4 °C overnight. Samples were rinsed three times in 0.1 M sodium cacodylate buffer (pH 7.2–7.4) and osmicated using 1% osmium tetroxide in cacodylate buffer. Following osmication, the samples were dehydrated using ascending ethanol concentrations, followed by two rinses in propylene oxide. Infiltration of the embedding medium was performed by immersion in a 1:1 mixture of propylene oxide and Epon (Science Services, Germany), followed by neat Epon and hardening at 60 °C for 48 h. For light microscopy, semi-thin sections (0.5 μm) of wellrounded and organized organoids were mounted on glass slides and stained for 1 min with 1% toluidine blue. For electron microscopy, ultra-thin sections (60 nm) were further cut and mounted on copper grids and stained using uranyl acetate and lead citrate. The sections were analysed with a JEM-2100Plus Transmission Electron Microscope at 200 kV (Jeol, Germany). Images were acquired with the XAROSA CMOS camera (Emsis, Germany).

### Western Blotting

Cells were lysed on ice with lysis buffer (10 mM Tris/HCl pH 7.4, containing 150 mM NaCl, 1% Triton X-100, 1x Protease Inhibitory Cocktail) for 30 min and centrifuged at 16,000 g for 15 min. Supernatants were collected and protein content quantified using BCA protein determination assay (Roth). Samples used for SDS-PAGE were added with SDS loading dye (500 mM Tris/HCl pH 6.8, 4% SDS, 40% glycerin, 40 mM DTT) and boiled at 95 °C for 5 minutes. The samples were subjected to SDS-PAGE, blotted onto nitrocellulose membranes (Amersham Protran). Membranes were blocked for 30 min with 25 mM Tris-buffered saline pH 7.4 (TBS) containing 5% milk powder and 0.05% Tween-20 and further incubated overnight at 4 °C with the following antibody dilutions in blocking buffer: rat anti Lamp1 (1:1000; clone 1D4B DHSB), rabbit anti CTSD, goat ant CTSZ (1:1000; R&D AF1033), rabbit anti GAPDH (1:1000; Santa Cruz Biotechnology sc-25778), rabbit anti Borcs5 (1:500; Proteintech 17169-1-AP) , rabbit anti BORCS7 (1:350, Biozol Abnova ABN-PAB23142), rabbit anti LC3-B (1:1,500; Novusbio NB100-2220), rabbit anti ARL8A (1:500, Proteintech 17060-1-AP). The membranes were then washed three times in 0.05% Tween-20 in TBS (TBS-T) and incubated 1 h at room temperature with the appropriate HRP-conjugated secondary antibody (Biozol). ECL detection was performed according to manufactureŕs instructions using Clarity substrate (Bio-Rad). Blots were imaged on ChemiDoc Universal Hodd II (Bio-Rad) using QuantityOne – 4.6.9 software and images prepared with ImageLab software 5.1.

### Proteomic and lipidomic analysis of magnetite-isolated lysosomes of MODE-K cells

Confluent WT and *Gnptab KO* MODE-K cells grown on 10 cm dishes for 48 h, were incubated in fresh medium supplemented with 10% v/v dextran-coated magnetite beads (DexoMAG40, Liquids Research Limited) for 14 h. After removal of magnetite medium and three PBS washing steps, the cells were further incubated in fresh magnetite-free culture medium for 12 h at 37 °C to allow the depletion of dextran beads in the endocytic pathway and their final delivery to lysosomes. Magnetite-loaded lysosomes were isolated from MODE-K cells as recently described [15]. For subsequent proteome and lipidome analyses three and five independent preparations, respectively, performed within 3 weeks were used.

### Data-independent acquisition (DIA) proteomics

#### Sample Preparation

SDS was added to the samples to a final concentration of 1% and samples were incubated at 95 °C for 10 min and subsequently homogenized using sonication using a Bioruptor® Homogenizer (Diagenode). Proteins were precipitated by addition of four sample volumes of ice-cold acetone, incubation over night at -20 °C, and centrifugation for 30 min at 20,000 x g and 4 °C. Precipitated proteins were washed once with 1 mL of ice-cold acetone and air-dried, proteins resuspended in 0.25 % RapiGest (Waters) and solubilized by incubation at 95 °C as well as sonication using a Bioruptor (Diagenode). Protein amounts were determined using the DC Protein Assay (Bio-Rad) and 10 g of protein from each sample were used for digestion following the SP3 protocol as described elsewhere [46] Briefly, proteins were reduced using a final concentration of 5 mM DTT for 45 min at 56 °C, alkylated with 20 mM acrylamide for 30 min at RT [47], and quenched through addition of 5 mM DTT. Samples were combined with SP3 beads (GE Healthcare, bead population A: 65151205050250; population B: 45152105050250) at a protein to beads ratio of 1:100 [w/w], 96% ethanol was added and the sample/beads mixture was incubated for 10 min at RT. Tubes were placed in a magnetic rack, supernatants discarded, the beads washed with 80% ethanol, resuspended in 50 mM TEAB, and digested using LysC (enzyme to protein ration of 1:100) for 3 h at 37 °C and trypsin (enzyme to protein ratio of 1:100) over night at 37 °C. The following day, supernatants were transferred to new tubes, the peptides desalted using STAGE tips [48] and dried in a vacuum centrifuge.

#### LC-MS/MS Analysis

Peptides were resuspended in 5% acetonitrile (ACN), 5% formic acid (FA) and 25% per sample were analyzed using a Dionex UltiMate 3000 nano-UHPLC coupled to an Orbitrap Fusion Lumos mass spectrometer (both Thermo Fischer Scientific). Analytical columns were produced in house as follows: spray tips were generated using a P-2000 laser puller (Sutter Instruments) from fused silica capillaries (360 µm outer, 100 µm inner diameter) and packed with 3 µm ReprosilPur AQ C18 particles (Dr. Maisch). Samples were loaded onto a 40 cm analytical column at a flow rate of 850 nl/min with 100 % solvent A (0.1% FA) and the peptides were separated with 120 min (data-independent acquisition (DIA)) or 240 min (data-dependent acquisition (DDA)) linear gradients from 3-35% solvent B (90% CAN, 0.1% FA) at a flow rate of 300 nL/min. For the DDA measurements, MS1 spectra were acquired at a mass range of m/z 350-1,200, an AGC target setting of 4×105, and a resolution of 60,000. Most intense precursor ions were fragmented in the top speed mode with an isolation width of m/z 1.6 and fragmentation by higher energy collisional dissociation (HCD, normalized collision energy (NCE) of 30%). MS2 spectra were acquired in the Orbitrap mass analyzer at a resolution of 30,000. Fragmented precursor ions were excluded from further fragmentation for 120 s. For DIA measurements, MS1 spectra were acquired at a mass range of m/z 350-1,200 in the Orbitrap mass analyzer with an AGC target setting of 5×105, a resolution of 120,000, and a maximum injection time of 20 msec. MS2 covered the MS1 scan range through 36 windows of 24.1 m/z each with an overlap of 0.5 m/z resulting in a cycle time of 3.44 sec. MS2 spectra were generated with 30% NCE and acquired in the Orbitrap mass analyzer at a resolution of 30,000, an AGC target setting of 1×106, and a maximum injection time of 60 msec.

#### MS Data Analysis

Thermo*.raw files were analyzed using Spectronaut (version 17.5.230413.5595). A hybrid DDA/DIA library was generated using the integrated Pulsar search engine and Uniprot Mus musculus (released 2024, 21,751 entries) and a database containing common contaminants (381 entries) [49]). For library generation and database searching default settings were applied. Trypsin/P was set as enzyme with a maximum of two missed cleavages, propionamide at cysteine was set as fixed modification and oxidation at methionine as variable modification; for library generation, the most abundant 3 – 6 fragment ions were selected automatically. The indexed-retention time concept (IRT, setting: dynamic) was applied for retention time alignment, and mass tolerance for MS1/MS2 ions as well as peak extraction windows were determined automatically by Spectronaut. Only precursor information was used for peak detection and global normalization (median-based) was performed. P-values were determined within the post-analysis pipeline of Spectronaut with default parameters.

### Targeted Mass Spectrometric Methods

#### Sample lysis and peptide preparation

Mouse organoids were stored at -80°C. For proteomic processing, they were thawed on ice and lysed in 4% SDS 0.1 M HEPES pH 7.5 by incubation at 95°C for 10 min, sonication using a BioRuptor Plus (Diagenode SA, Belgium) for 20 cycles (15 sec on/off, setting: high), and additional incubation at 95°C for 10 min. Lysates were cleared by centrifugation (16,000 xg, 30 min), protein amounts were determined using the DC protein assay (BioRad, Hercules, CA), and 100 µg of each sample subjected to SP3 protein digestion as described elsewhere [46] Briefly, protein lysates were reduced with 20 mM DTT (30 min, 56 °C), alkylated with 40 mM acrylamide (30 min, room temperature) and quenched with 20 mM DTT (15 min, room temperature). Afterwards, SP3-bead mixture was added at a 10:1 bead-to-protein ratio, binding induced by adding 96% ethanol, samples placed on a magnet to collect beads, and supernatants discarded. Beads were washed in 80% ethanol, the supernatant discarded, and they were resuspended in 50 mM TEAB, pH 8.0 containing proteomics grade trypsin (Promega, Madison, WI) at a 1:25 protease-to-protein ratio. Samples were digested for 18 h at 37 °C, 1000 rpm, the supernatants containing peptides transferred to a new microtube, acidified by acetic acid (2% final concentration), and desalted using self-packed StageTips [48] containing Empore C18 material (CDS Analytical, Oxford, PA). The cleaned peptide extracts were dried in a vacuum centrifuge, dissolved in 5% acetonitrile, peptide amounts determined using the Pierce Fluorometric Peptide Assay kit (Thermo Fisher Scientific) and samples stored at -20°C.

#### LC-MS analysis

Dissolved peptides were mixed with indexed retention time (iRT) standard peptides [50] dried in a vacuum centrifuge and redissolved in 5% ACN, 5% formic acid (FA). For LC-MS/MS analysis, 1 ug of peptide were separated on a Dionex Ultimate 3000 RSLC nano HPLC system (Dionex, Idstein, Germany) coupled to an Orbitrap Fusion Lumos mass spectrometer (Thermo Fisher Scientific, Bremen, Germany). Peptides were injected onto a C18 analytical column (400 mm length, 100 µm inner diameter, ReproSil-Pur 120 C18-AQ, 3 µm, made in-house), which was equilibrated at 99% solvent A (0.1% FA) and 1% solvent B (90% ACN, 0.1% FA). For targeted analyses, 596 peptides covering 321 mouse lysosomal proteins were measures by a previously established parallel reaction monitoring (PRM) assay [42] using 8 min scheduling windows. Peptides were separated with a 120 min linear gradient from 1 to 35 % solvent B and MS1 spectra acquired from 300 to 1500 m/z every 3 seconds in the Orbitrap analyzer at a resolution of 60,000, a maximum injection time of 118 ms, and an automatic gain control target setting of 4e5. Lock mass correction was enabled. Target ions were subjected to HCD fragmentation (normalized collision energy 27%), and product ions analyzed from 200 - 2000 m/z in the Orbitrap analyzer with a resolution of 30,000, a maximum injection time of 54 ms, and an automatic gain control target of 50,000. Isolation lists were generated based on an initial scheduling run in Skyline-daily [51] (version 21.1.1.160) with the iRT calculator.

#### Data analysis

Data from targeted measurements were analyzed in Skyline-daily. Validation of MS2 spectra and peak picking was aided by a sample specific spectral library created by searching the PRM raw files with Proteome Discoverer (version 2.5.0.400, Thermo Fisher Scientific, Bremen, Germany) using Mascot (version 2.6.1, Matrix Science, London, UK). Data were searched against SwissProt including isoforms with species specificity set to “Mus musculus” in combination with the common contaminants databases cRAP (https://www.thegpm.org/crap/) and MaxQuant contaminants. Trypsin/P was selected as enzyme, two missed cleavages were allowed, and mass tolerances set to 20/40 ppm for MS1/MS2 spectra, respectively. Propionamide (C) was defined as fixed modification, and Oxidation (M) as well as Acetyl (Protein N-term) as variable modifications. Identifications were validated by Percolator [52] (version 3.05.0) and the resulting msf file was loaded into Skyline to build the spectral library. Peptide level MS2-Quantification was performed in Skyline based on summed peak areas of the 3-6 most intense fragment ions (type: b, y, charge: 1+, 2+) and peptide peak areas together with the corresponding library dot product (dotp) values were exported as txt file. Further data analysis was performed using Perseus [53] (version 1.6.15.0). First, peak area values were log2-transformed, followed by removal of peptides with dotp values ≤ 0.6 and peptides quantified in only one sample. MLII/WT peptide ratios were calculated for all possible replicate combinations followed by protein ratio calculation using the median of all peptides belonging to the same protein. Finally, differentially expressed proteins were determined based on a one-sample t-test against a fixed value of 0. Proteins exhibiting an absolute log2 fold change ≥ 0.58 and a q-value ≤ 0.05 (Benjamini-Hochberg method) were deemed differentially expressed. Volcano plots were generated in R (version 4.3, https://cran.r-project.org) utilizing base and the tidyverse (version 2.0) packages.

### Lipidomics analysis

#### Lipid extraction

120 µL ice cold MeOH containing a mix of deuterated standards were added to the magnetite-purified lysosomal fractions, incubated at -30°C for 1 min and, afterwards, put in a sonic bath for 10 min. After adding 400 µL of ice-cold MTBE samples were re-incubated in a Thermomixer for 1h × 4°C × 1200 rpm followed by addition of 188 µL of water and centrifugation, upper phases were collected and dried. The extracts were resuspended in 100 µL of CHCl_3_/MeOH/IPA (1/2/4 *v/v*) and analyzed by LC-MS/MS.

#### RP-UHPLC-TIMS-MS method parameters

UHPLC-TIMS-MS analyses were performed on an Ultimate RS3000 UHPLC (Thermo Fisher Scientific, Milan, Italy). The LC system was coupled online to a TimsTOF Pro Quadrupole Time of Flight (Q-TOF) (Bruker Daltonics, Bremen, Germany) equipped with an Apollo II electrospray ionization (ESI) probe. Lipid separation was performed with an Acquity UPLC CSH^TM^ C18 column (100 × 2.1 mm; 1.7 μm, 130 Å) protected with a VanGuard CSHTM precolumn (50 × 2.1 mm; 1.7 μm, 130 Å) (Waters, Milford, MA, U.S.A). Column temperature was set to 65 °C, flow rate was set to 0.5 mL min^-1^, mobile phase consisted of (A): ACN/H_2_O 60:40 (*v/v %*) and (B): IPA/ACN 90:10 (*v/v %*) both buffered with 10 mM HCOONH_4_ and 0.1% HCOOH (*v/v %*). The following gradient has been used: 0 min, 50% B; 0.6 min, 50% B; 2.0 min, 52% B; 5.0 min, 58% B; 6.5 min, 80% B; 6.6 min, 95% B; 8.6 min, 99% B; 8.7 min, 50% B and then 1.3 min for column re-equilibration. Prior to each LC-MS run, the TimsTOF Pro was recalibrated in mass and mobility with a mixture (*1:1 v/v %*) of 10 mM sodium formate calibrant solution and ESI-L Low Concentration Tuning Mix. The TIMS-MS analyses were performed in data-dependent parallel accumulation serial fragmentation (DDA-PASEF) in both positive and negative ionization, in separate runs. The injection volume was set at 2 µL for ESI^+^ and at 5 µL for ESI^-^. Source parameters: Nebulizer gas (N_2_) pressure: 4.0 Bar, Dry gas (N_2_): 10 L/min, Dry temperature: 280°C. Mass spectra were recorded in the range m/z 50–1500, with an accumulation and ramp time to 100 ms each. The ion mobility was scanned from 0.55 to 1.80 Vs/cm^2^. Precursors for data-dependent acquisition were isolated within ± 2 m/z and fragmented with a TIMS-STEPPING ion mobility-dependent collision energy mode: CE [eV] #1: 20-40 and CE [eV] #2: 35-50. Exclusion time was set to 0.1 min, Ion charge control (ICC) was set to 7.5 Mio.

#### RP-UHPLC-TIMS-MS data analysis and processing

4D data alignment, filtering and annotation were performed with MetaboScape 2023b (Bruker) employing a feature finding algorithm (T-Rex 4D) that automatically extracts buckets from raw files. Feature detection was set to 250 and 150 counts for positive and negative modes. The minimum number of data points in the 4D-TIMS space was set to 100 and recursive feature extraction was used (75 points).

#### Lipid annotation

Lipid annotation was performed first with a rule-based annotation, based on diagnostic-class specific fragments and their intensity in acquired MS/MS spectra, and, subsequently, using the LipidBlast spectral library of MS DIAL (http://prime.psc.riken.jp/compms/msdial/main.html) with the following parameters: Mass accuracy: narrow 2 ppm, wide 10 ppm; mSigma: narrow 30, wide 250, MS/MS score: narrow 800, wide 150. Collision cross-section (CCS) %: narrow 1, wide 3.5. The spectra were processed in positive mode using [M+H]^+^, [M+Na] ^+^, [M+K] ^+^, [M+H–H_2_O] ^+^ and [M+NH_4_] ^+^ ions, while [M–H]^−^, [M+Cl] ^−^, [M+HCOO] ^−^ and [M–H_2_O] ^−^ in negative mode. The assignment of the molecular formula was performed for the detected features using Smart Formula™ (SF). Each lipid feature was manually curated following Lipidomics Standard Initiative (LSI) guidelines (https://lipidomics-standards-initiative.org/guidelines/lipid-species-identification/general-rules). Additionally, LipidCreator tool (https://lifs-tools.org/lipidcreator.html) extension in Skyline (https://skyline.ms/project/home/begin.view) was used for in silico comparison of specific product ions for manual MS/MS curation.

### Measurement of lysosomal pH

Lysosomal pH was measured as previously described [54]. MODE-K cells were plated on glass-bottom live-cell dishes (MatTek) 24 h before pH measurement. Cells were loaded with 0.5 mg/ml Oregon Green 488-dextran (Life Technologies GmbH) in growth medium overnight, followed by a 2 h chase for complete dye delivery to lysosomes. For imaging, the medium was changed to an imaging buffer at pH 7.4. Images were acquired using a Leica Dmi8 Thunder Imager microscope (Leica Microsystems) equipped with a 63x 1.40 NA oil-immersion lens and an Oregon Green filter cube (AHF) with excitation at 440 nm or 480 nm, respectively. In situ pH calibration curves were subsequently obtained for each dish, using isotonic K-based buffers supplemented with 10 μM nigericin and 10 μM monensin after equilibration for at least 2 min for each pH (ranging between pH 3.5 - 6.0) starting with pH 6. Images were analyzed using ImageJ, where regions of interest (ROIs) were defined as areas above a defined fluorescence threshold in the acquired images at 440-nm excitation. The mean intensity ratio between 480- (pH-sensitive) and 440-nm excitation (pH-insensitive) was calculated after background subtraction for each ROI. The fluorescence intensity ratios (480/440) as a function of pH were fit to a sigmoid and used to interpolate the pH values from the cells.

### Autophagic flux measurements

24h after seeding MODE-K cells on MatTek glass bottom dishes, cells were transiently transfected with tandem RFP-GFP-LC3 (ptfLC3 [32]) using Lipofectamine 2000. After 24 h, growth medium was replaced by prewarmed imaging buffer (135 mM NaCl, 5 mM KCl, 2 mM CaCl_2_, 1 mM MgCl_2_, 10 mM HEPES pH 7.4, 10 mM glucose) and confocal images acquired using an 63X oil immersion objective on a LSM880 confocal microscope equipped with an Airyscan detector (Zeiss). Images were processed using ImageJ. A binary mask was generated based on the RFP fluorescence image, applied to both images, and the GFP:RFP ratio calculated in Excel. ptfLC3 was a gift from Tamotsu Yoshimori (Addgene plasmid # 21074 ; http://n2t.net/addgene:21074 RRID:Addgene_21074)

## Acknowledgements

The authors would like to thank Emanuela Szpotowicz (ZMNH, Hamburg) for expert technical assistance. Philip Rosenstiel (IKMB, Kiel) and Stephan Storch (UKE, Hamburg) kindly provided reagents.

## Author contributions

Gianmarco del Gallo: Investigation, Methodology, Formal analysis; Zilei Chen: Investigation, Methodology; Anne Sanner: Methodology Formal analysis (Proteomics); Robert Hardt: Methodology, Formal analysis (Proteomics); Fabrizio Merciai Methodology, Formal analysis (Lipidomics); Fabiola Salsano: Investigation, Shroddha Bose: Methodology, Formal analysis (pH measurement); Michaela Schweizer Investigation, Methodology (EM); Diego Medina: Conceptualization (Lipidomics); Tobias Stauber: Methodology (pH measurement), Supervision, Writing – review and editing; Jens Bosse: Methodology (LLS), Supervision; Eduardo M. Sommella, Formal analysis, methodology (Lipidomics), Supervision; Dominic Winter: Methodology (Proteomics), Supervision; Sabrina Jabs: Investigation, Formal analysis, Supervision, Resources, Writing – original draft; Thomas Braulke: Conceptualization, Supervision, Resources, Writing – original draft.

## Data availability statement

All primary data and resources included in this study are available from the corresponding authors upon request.

## Abbreviations

ASM: acid sphingomyelinase (*Smpd1*)
BMP: bis(monoacylglycero)phosphate
Bodipy-LacCer: Bodipy-labeled lactosylceramide
Cer: ceramide
CTSD: cathepsin D
GNPTAB: GlcNAc-1-phosphotransferase
IP: immunoprecipitation
KI: knock-in
KO: knock-out
LPC: lysophosphatidylcholine
M6P: mannose 6-phosphate
MODE-K: mouse duodenal epithelial cell clone K
MS: mass spectrometry
NPC2: Niemann-Pick C disease cholesterol transporter 2
PLA2G15: phospholipase A2 group 15
PSAP: prosaposin activator protein
SD: standard deviation
SM: sphingomyelin
SQSTM1: sequestosome 1
WT: wild-type

---

**Fig. S1.**
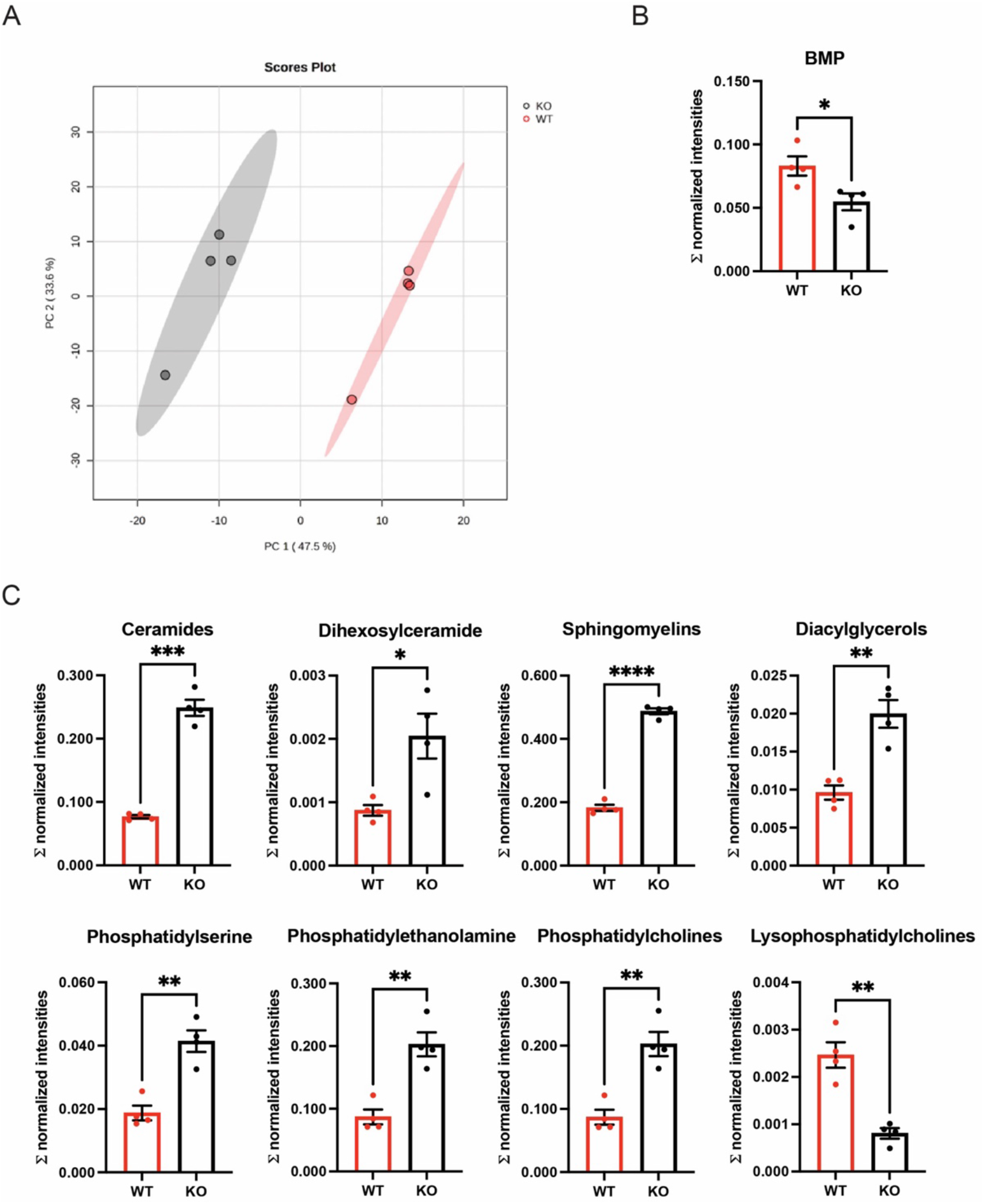
Analysis of differentially expressed lipids in WT and *Gnptab* KO MODE-K cells. (**A**) Principal component analysis (PCA) plot showing the strong differences between the lysosomal lipids color-coded for the two genotype groups (PC1). (**B,C**) Significantly altered lipid subclasses quantified as sum of normalized intensities (mean ± SD; unpaired student t-test with Welch’s correction; n=4 of each genotype) \**p*< 0.05; ** *p*<0.01; *** *p*<0.001; **** *p*<0.0001

**Figure S2.**
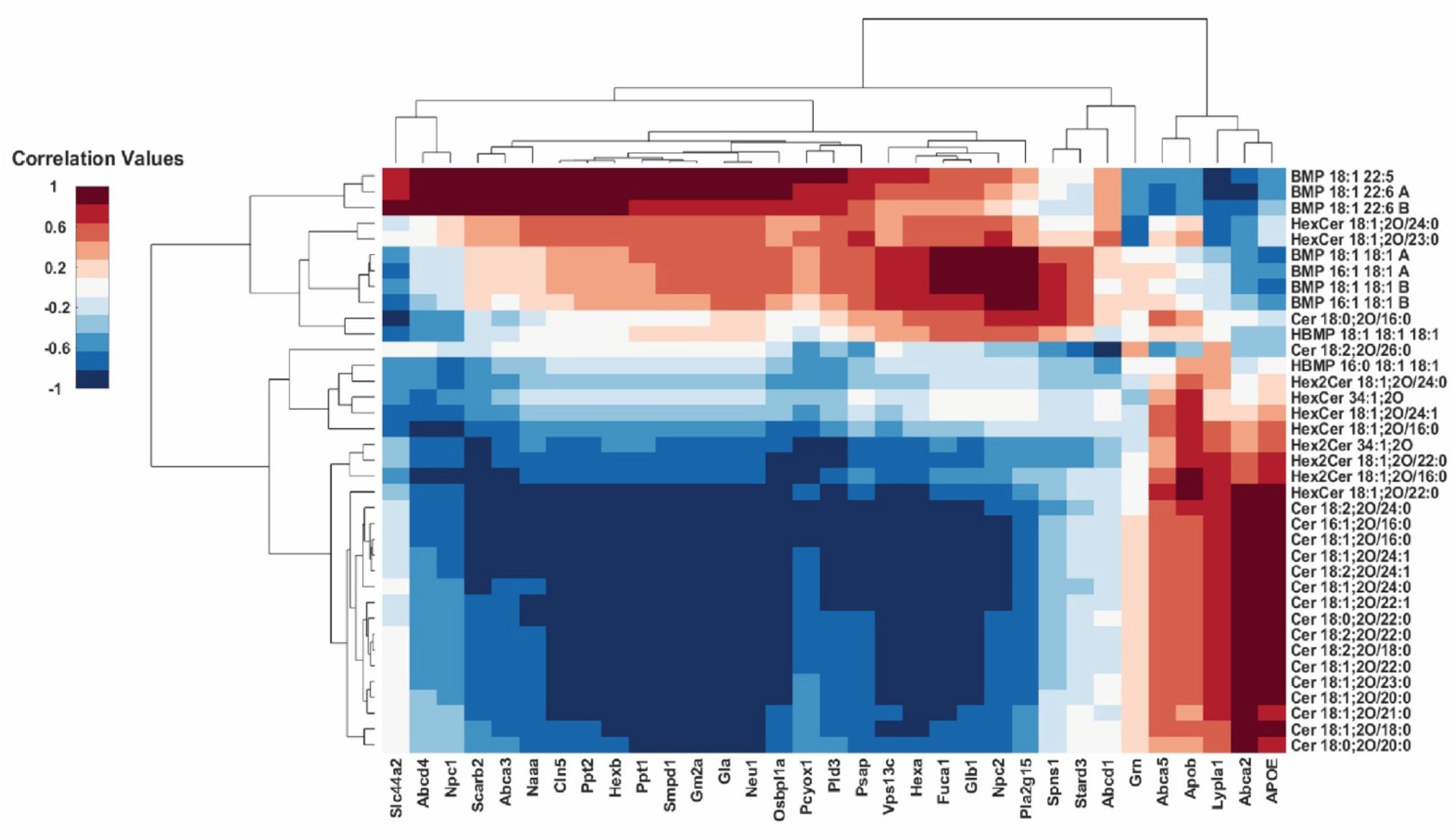
Correlation of the lysosomal lipidome and lysosomal proteins involved in lipid metabolism and egress. Letters “A” and “B” indicate possible isomers.

**Figure S3.**
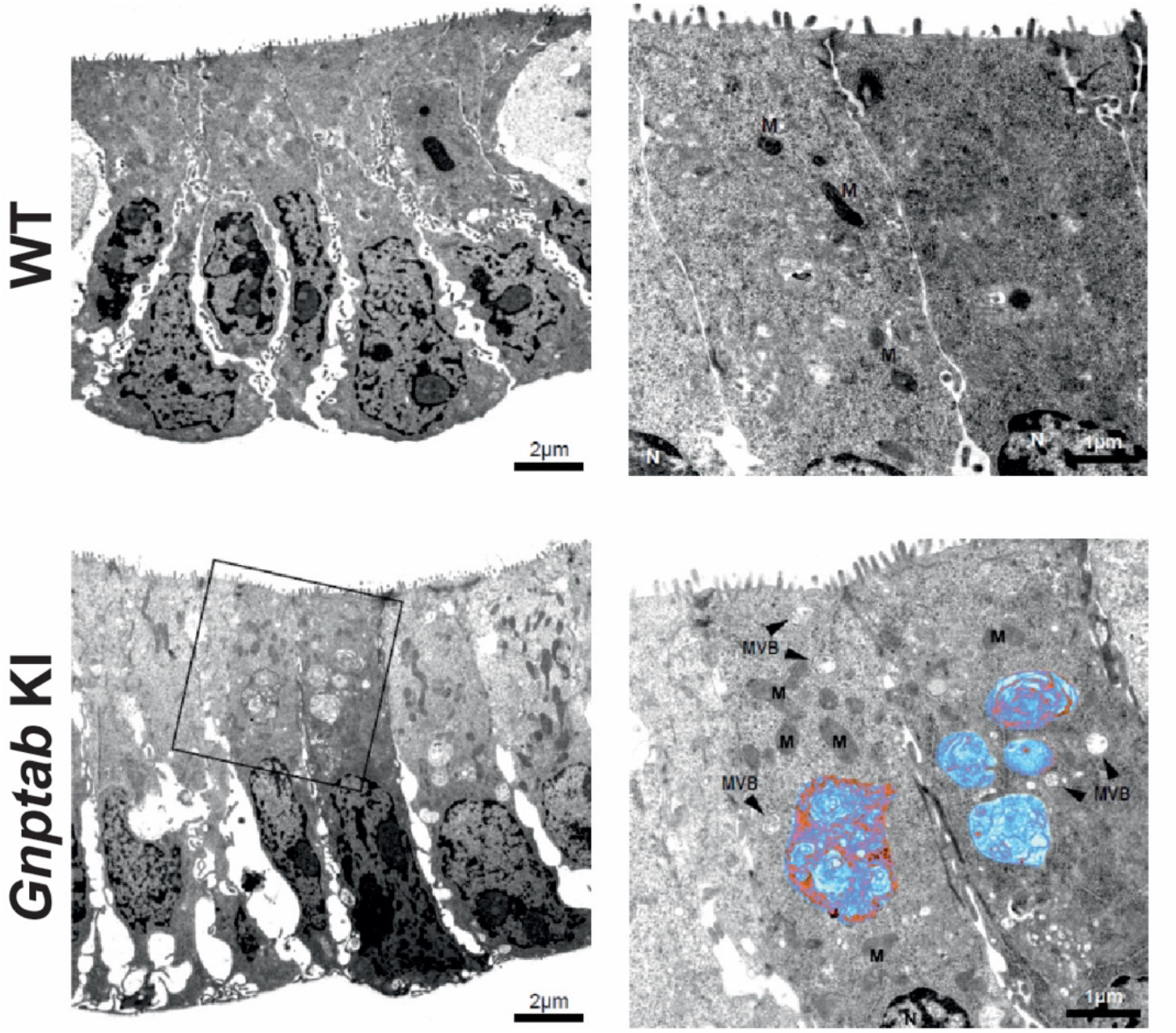
Electron Microscopy images of thin sections of resin-embedded WT and *Gnptab*-KI organoids. In enterocytes of WT organoids (upper images) no lysosomes were detectable (magnified image). *Gnptab* KI organoids (lower images) showed several enlarged lysosomes (marked in blue; magnified image). M=mitochondria, MVB=multivesicular bodies, N=nuclei.

## References

[1] Braulke T, Carette JE, Palm W. Lysosomal enzyme trafficking: from molecular mechanisms to human diseases. Trends in Cell Biology. 2024 2024/03/01/;34(3):198–210.

[2] Rudnik S, Damme M. The lysosomal membrane—export of metabolites and beyond. The FEBS Journal. 2021 2021/07/01;288(14):4168–4182.

[3] Ballabio A, Bonifacino JS. Lysosomes as dynamic regulators of cell and organismal homeostasis. Nature Reviews Molecular Cell Biology. 2020 2020/02/01;21(2):101–118.

[4] Chadwick SR, Grinstein S, Freeman SA. From the inside out: Ion fluxes at the centre of endocytic traffic. Current Opinion in Cell Biology. 2021 2021/08/01/;71:77–86.

[5] Settembre C, Perera RM. Lysosomes as coordinators of cellular catabolism, metabolic signalling and organ physiology. Nature Reviews Molecular Cell Biology. 2024 2024/03/01;25(3):223–245.

[6] Tiede S, Storch S, Lübke T, et al. Mucolipidosis II is caused by mutations in GNPTA encoding the α/β GlcNAc-1-phosphotransferase. Nature Medicine. 2005 2005/10/01;11(10):1109–1112.

[7] Cathey SS, Leroy JG, Wood T, et al. Phenotype and genotype in mucolipidoses II and III alpha/beta: a study of 61 probands. Journal of Medical Genetics. 2010;47(1):38.

[8] Hickman S, Neufeld EF. A hypothesis for I-cell disease: Defective hydrolases that do not enter lysosomes. Biochemical and Biophysical Research Communications. 1972 1972/11/15/;49(4):992–999.

[9] Markmann S, Krambeck S, Hughes CJ, et al. Quantitative Proteome Analysis of Mouse Liver Lysosomes Provides Evidence for Mannose 6-phosphate-independent Targeting Mechanisms of Acid Hydrolases in Mucolipidosis II*. Molecular & Cellular Proteomics. 2017 2017/03/01/;16(3):438–450.

[10] Markmann S, Thelen M, Cornils K, et al. Lrp1/LDL Receptor Play Critical Roles in Mannose 6-Phosphate-Independent Lysosomal Enzyme Targeting. Traffic. 2015;16(7):743–759.

[11] Lefrancois S, Zeng J, Hassan AJ, et al. The lysosomal trafficking of sphingolipid activator proteins (SAPs) is mediated by sortilin. The EMBO Journal. 2003 2003/12/01;22(24):6430–6437.

[12] Vidal K, Grosjean I, Revillard J-P, et al. Immortalization of mouse intestinal epithelial cells by the SV40-large T gene: Phenotypic and immune characterization of the MODE-K cell line. Journal of Immunological Methods. 1993 1993/11/05/;166(1):63–73.

[13] Kollmann K, Damme M, Markmann S, et al. Lysosomal dysfunction causes neurodegeneration in mucolipidosis II ‘knock-in’ mice. Brain. 2012;135(9):2661–2675.

[14] Kraus F, He Y, Swarup S, et al. Global cellular proteo-lipidomic profiling of diverse lysosomal storage disease mutants using nMOST. Science Advances. 2025;11(4):eadu5787.

[15] Richards CM, Jabs S, Qiao W, et al. The human disease gene LYSET is essential for lysosomal enzyme transport and viral infection. Science. 2022;378(6615):eabn5648.

[16] Pechincha C, Groessl S, Kalis R, et al. Lysosomal enzyme trafficking factor LYSET enables nutritional usage of extracellular proteins. Science. 2022;378(6615):eabn5637.

[17] Hasilik A, Neufeld EF. Biosynthesis of lysosomal enzymes in fibroblasts. Phosphorylation of mannose residues. J Biol Chem. 1980 May 25;255(10):4946–50.

[18] Lee WS, Payne BJ, Gelfman CM, et al. Murine UDP-GlcNAc:lysosomal enzyme N-acetylglucosamine-1-phosphotransferase lacking the gamma-subunit retains substantial activity toward acid hydrolases. J Biol Chem. 2007 Sep 14;282(37):27198–27203.

[19] De Pace R, Ghosh S, Williamson CD, et al. BLOC-1 and BORC: Complex regulators of endolysosomal dynamics. Cell Chemical Biology. 2025;32(9):1106–1124.

[20] Roney JC, Li S, Farfel-Becker T, et al. Lipid-mediated impairment of axonal lysosome transport contributing to autophagic stress. Autophagy. 2021 2021/07/03;17(7):1796–1798.

[21] Roney JC, Li S, Farfel-Becker T, et al. Lipid-mediated motor-adaptor sequestration impairs axonal lysosome delivery leading to autophagic stress and dystrophy in Niemann-Pick type C. Developmental Cell. 2021;56(10):1452–1468.e8.

[22] Ebner M, Fröhlich F, Haucke V. Mechanisms and functions of lysosomal lipid homeostasis. Cell Chemical Biology. 2025;32(3):392–407.

[23] Weesner JA, Annunziata I, van de Vlekkert D, et al. Glycosphingolipids within membrane contact sites influence their function as signaling hubs in neurodegenerative diseases. FEBS Open Bio. 2023 2023/09/01;13(9):1587–1600.

[24] Medoh UN, Hims A, Chen JY, et al. The Batten disease gene product CLN5 is the lysosomal bis(monoacylglycero)phosphate synthase. Science. 2023 2023/09/15;381(6663):1182–1189.

[25] Medoh UN, Abu-Remaileh M. The Bis(monoacylglycero)-phosphate Hypothesis: From Lysosomal Function to Therapeutic Avenues. Annu Rev Biochem. 2024 Aug;93(1):447–469.

[26] Oninla VO, Breiden B, Babalola JO, et al. Acid sphingomyelinase activity is regulated by membrane lipids and facilitates cholesterol transfer by NPC2 [S]. Journal of Lipid Research. 2014;55(12):2606–2619.

[27] Abdul-Hammed M, Breiden B, Adebayo MA, et al. Role of endosomal membrane lipids and NPC2 in cholesterol transfer and membrane fusion [S]. Journal of Lipid Research. 2010;51(7):1747–1760.

[28] Makrypidi G, Damme M, Müller-Lönnies S, et al. Mannose 6 Dephosphorylation of lysosomal proteines mediated by acid phosphatases Acp2 and Acp5. Molecular and Cellular Biology. 2012;32:774–782.

[29] Gabandé-Rodríguez E, Boya P, Labrador V, et al. High sphingomyelin levels induce lysosomal damage and autophagy dysfunction in Niemann Pick disease type A. Cell Death & Differentiation. 2014 2014/06/01;21(6):864–875.

[30] Hodul M, Lane-Donovan C, Cheang ES, et al. Prosaposin Is Cleaved Into Saposins by Multiple Cathepsins in a Progranulin-Regulated Fashion. Journal of Neurochemistry. 2026 2026/01/01;170(1):e70357.

[31] Nixon RA, Rubinsztein DC. Mechanisms of autophagy–lysosome dysfunction in neurodegenerative diseases. Nature Reviews Molecular Cell Biology. 2024 2024/11/01;25(11):926–946.

[32] Kimura S, Noda T, Yoshimori T. Dissection of the Autophagosome Maturation Process by a Novel Reporter Protein, Tandem Fluorescent-Tagged LC3. Autophagy. 2007 2007/09/20;3(5):452–460.

[33] Yan J, Zhang Y, Choksi S, et al. TGFB-inducible VASN (vasorin) promotes lysosomal acidification. Autophagy. 2026 2026/05/04;22(5):1097–1115.

[34] Huang H, Ouyang Q, Zhu M, et al. mTOR-mediated phosphorylation of VAMP8 and SCFD1 regulates autophagosome maturation. Nature Communications. 2021 2021/11/16;12(1):6622.

[35] Vargas JNS, Hamasaki M, Kawabata T, et al. The mechanisms and roles of selective autophagy in mammals. Nature Reviews Molecular Cell Biology. 2023 2023/03/01;24(3):167–185.

[36] Yamamoto H, Zhang S, Mizushima N. Autophagy genes in biology and disease. Nature Reviews Genetics. 2023 2023/06/01;24(6):382–400.

[37] Xue Q, Kang R, Klionsky DJ, et al. Copper metabolism in cell death and autophagy. Autophagy. 2023 2023/08/03;19(8):2175–2195.

[38] Klionsky DJ, Abdel-Aziz AK, Abdelfatah S, et al. Guidelines for the use and interpretation of assays for monitoring autophagy (4th edition)1. Autophagy. 2021 2021/01/02;17(1):1–382.

[39] Kollmann K, Damme M, Markmann S, et al. Lysosomal dysfunction causes neurodegeneration in mucolipidosis II ’knock-in’ mice. Brain. 2012 Sep;135(Pt 9):2661–75.

[40] Kollmann K, Pestka JM, Kühn SC, et al. Decreased bone formation and increased osteoclastogenesis cause bone loss in mucolipidosis II. EMBO Molecular Medicine. 2013;5(12):1871–1886.

[41] Otomo T, Schweizer M, Kollmann K, et al. Mannose 6 phosphorylation of lysosomal enzymes controls B cell functions. Journal of Cell Biology. 2015;208(2):171–180.

[42] Mosen P, Sanner A, Singh J, et al. Targeted Quantification of the Lysosomal Proteome in Complex Samples. Proteomes. 2021;9(1):4.

[43] Sly WS, Vogler C, Grubb JH, et al. Enzyme therapy in mannose receptor-null mucopolysaccharidosis VII mice defines roles for the mannose 6-phosphate and mannose receptors. Proceedings of the National Academy of Sciences. 2006 2006/10/10;103(41):15172–15177.

[44] Ootani A, Li X, Sangiorgi E, et al. Sustained in vitro intestinal epithelial culture within a Wnt-dependent stem cell niche. Nature Medicine. 2009 2009/06/01;15(6):701–706.

[45] Mahe MM, Aihara E, Schumacher MA, et al. Establishment of Gastrointestinal Epithelial Organoids. Current Protocols in Mouse Biology. 2013 2013/12/01;3(4):217–240.

[46] Bonini S, Winter D. Two-Step Enrichment Facilitates Background Reduction for Proteomic Analysis of Lysosomes. Journal of Proteome Research. 2024;23(8):3393–3403.

[47] Müller T, Winter D. Systematic Evaluation of Protein Reduction and Alkylation Reveals Massive Unspecific Side Effects by Iodine-containing Reagents*. Molecular & Cellular Proteomics. 2017 2017/07/01/;16(7):1173–1187.

[48] Rappsilber J, Ishihama Y, Mann M. Stop and Go Extraction Tips for Matrix-Assisted Laser Desorption/Ionization, Nanoelectrospray, and LC/MS Sample Pretreatment in Proteomics. Analytical Chemistry. 2003;75(3):663–670.

[49] Frankenfield AM, Ni J, Ahmed M, et al. Protein Contaminants Matter: Building Universal Protein Contaminant Libraries for DDA and DIA Proteomics. Journal of Proteome Research. 2022;21(9):2104–2113.

[50] Escher C, Reiter L, MacLean B, et al. Using iRT, a normalized retention time for more targeted measurement of peptides. PROTEOMICS. 2012 2012/04/01;12(8):1111–1121.

[51] Pino LK, Searle BC, Bollinger JG, et al. The Skyline ecosystem: Informatics for quantitative mass spectrometry proteomics. Mass Spectrometry Reviews. 2020 2020/05/01;39(3):229–244.

[52] Käll L, Canterbury JD, Weston J, et al. Semi-supervised learning for peptide identification from shotgun proteomics datasets. Nature Methods. 2007 2007/11/01;4(11):923–925.

[53] Tyanova S, Temu T, Sinitcyn P, et al. The Perseus computational platform for comprehensive analysis of (prote)omics data. Nature Methods. 2016 2016/09/01;13(9):731–740.

[54] Weinert S, Jabs S, Supanchart C, et al. Lysosomal pathology and osteopetrosis upon loss of H+-driven lysosomal Cl-accumulation. Science. 2010 Jun 11;328(5984):1401–3.

